# Loss of Progranulin Results in Increased Pan-Cathepsin Activity and Reduced LAMP1 Lysosomal Protein

**DOI:** 10.1101/2023.07.15.549151

**Authors:** Abigail Anderson, Malú G. Tansey

## Abstract

Mutations in the progranulin (PGRN) encoding gene, *GRN*, cause familial frontotemporal dementia (FTD) and neuronal ceroid lipofuscinosis (NCL) and PGRN is also implicated in Parkinson’s disease (PD). These mutations result in decreased PGRN expression. PGRN is highly expressed in peripheral immune cells and microglia and regulates cell growth, survival, repair, and inflammation. When PGRN is lost, the lysosome becomes dysfunctional, but the exact mechanism by which PGRN plays a role in lysosome function and how this contributes to inflammation and degeneration is not entirely understood. To better understand the role of PGRN in regulating lysosome function, this study examined how loss of *GRN* impacts total LAMP1 protein expression and cathepsin activities. Using mouse embryonic fibroblasts (MEFs), immunocytochemistry and immunoblotting assays were performed to analyze fluorescent signal from LAMP1 (lysosomal marker) and BMV109 (marker for pan-cathepsin activity). *GRN*^*-/-*^ MEFs exhibit increased expression of pan-cathepsin activity relative to *GRN*^*+/+*^ MEFs, and significantly impacts expression of LAMP1. The significant increase in pan-cathepsin activity in the *GRN*^*-/-*^ MEFs confirms that PGRN loss does alter cathepsin expression, which may be a result of compensatory mechanisms happening within the cell. Using NTAP PGRN added to *GRN*^*-/-*^ MEFs, specific cathepsin activity is rescued. Further investigations should include assessing LAMP1 and BMV109 expression in microglia from *GRN*^*-/-*^ mice, in the hopes of understanding the role of PGRN in lysosomal function in immune cells of the central nervous system and the diseases in which it is implicated.

## Introduction

Progranulin (PGRN) protein is a secreted glycoprotein that can be processed into granulin peptides in extracellular space and uses multiple pathways to travel into the lysosome of cells (Kao et al., 2017). It is composed of seven subunits called granulins which can be cleaved by sortilin and prosaposin (Houser et al., 2022). PGRN protein is encoded by the *GRN* gene and mutations in this gene can result in either an under expression or overexpression of PGRN. Under expression of PGRN protein is associated with frontotemporal dementia (FTD) and neurodegeneration and overexpression of PGRN protein has been shown to worsen the pathology of neurodegeneration (Nguyen et al., 2018; Tanaka et al., 2022). PGRN protein is important in cell growth, survival and repair and it is known to be localized to the lysosome and highly expressed in multiple different cell types including peripheral immune cells and microglia (Nguyen et al., 2018).

PGRN protein has a working relationship with cathepsins exemplified by the way in which cathepsins help cleave PGRN protein into individual granulins and PGRN impacts cathepsin expression. (Simon et al., 2022). Cathepsins are proteases found within lysosomes that are associated with inflammatory neurodegenerative diseases (Yadati et al., 2020). Cathepsin activity can be measured by an *in vitro* assay with BMV109, a membrane permeable and fixable fluorescent pan-cathepsin probe (Staderini et al., 2018).

It is well established that lysosomes are instrumental in detecting the onset of infection and facilitating an immune response (Inpanathan & Botelho, 2019). For this study, an important immune response that the lysosome helps to facilitate is inflammation. Inflammation is an important immune response to consider when studying neurodegeneration because chronic inflammation develops as a function of age, a process that has been coined with the term “inflammaging” (Tansey et al., 2022). Keeping in mind that PGRN protein is localized to the lysosome (Nguyen et al., 2018), there is this relationship between the lysosome, inflammation, and neurodegeneration. Because of this, PGRN protein acts as a lysosome-related risk factor in multiple neurodegenerative diseases, with inflammation as a hallmark, including FTD, NCL and PD (Kao et al., 2017; Tansey et al., 2022; Zhou et al., 2017).

Because of the impact that PGRN protein has on inflammation and neurodegeneration (Eriksen & Mackenzie, 2008), there is a need to better understand this complex protein. If it is correct that PGRN protein is critical for lysosomal health, I would predict that cells without PGRN protein would have less lysosomal protein expression but more cathepsin activity. The aim of this study is to investigate the role of PGRN protein in lysosomal health and how this contributes to inflammation and neurodegeneration. Mouse embryonic fibroblasts (MEFs) are spindle-shaped cells harvested from a mouse embryo which make a stable cell line with a rapid growth rate which serves as a useful model to understand biochemical processes. In order to do this, *Grn+/+* MEFs and *Grn* -/- MEFs were used as tools to study the lysosome and other pathways affected by PGRN mutations.

## Methods

### Mouse Embryonic Fibroblasts (MEFs)

MEFs were cultured in complete media consisting of Dulbecco’s Modified Eagle Medium (DMEM) High Glucose (Gibco 11960-044) with 10% HI-FBS (R&D Systems S11150), 100x Pen/Strep (Gibco 15140-122), 100 mM sodium pyruvate (Gibco 11360-070) and GlutaMAX (100X) (Gibco 35050-061). MEFs were maintained by serial passing every 3-4 days. The *Grn*^*-/-*^ MEFs were generated in the lab of Dr. Robert Farese (*The Farese & Walther Lab* | *Sloan Kettering Institute*, n.d.).

### *In Vitro* Cathepsin Activity Assay using the BMV109 Fluorescent Probe

MEFs were plated in a 12-well plate at a density of 150,000 cells per well and incubated overnight at 37° C. The next day, two hours prior to adding the BMV109 (Vergent Bioscience 40200-200) probe, selected wells were treated with a final concentration of 40 nM Bafilomycin A1(Fermentek 88899-55-2) in complete media to inhibit autophagosome-lysosome fusion in control conditions. The probe was administered at a final concentration of 1 _μ_M in media for experimental conditions and a final concentration of 1 _μ_M BMV109 and 40 nM Bafilomycin A1 in control conditions and cells were incubated for one hour. Proteins were resolved by Mini-Protean TGX stain-free gels 4-20% (BioRad 4568093) and transferred to TransBlot Turbo mini-size PVDF membrane (BioRad). Membranes were fixed with 0.4% PFA then rinsed three times with water. BMV109 signal was detected using the Li-Cor Odyssey Fc system.

### PGRN Rescue by Transfection of N-TAP Progranulin

*E. coli* stock was grown up in lysogeny broth (LB) to confirm antibiotic resistance, then individual colonies of E. coli were cultured on LB agar plates. Individual colonies or clones were grown in LB broth with kanamycin. Plasmid was isolated using QIAprep Spin Miniprep Kit (ID #27104) according to its included protocol. MEFs were transfected with the isolated N-TAP progranulin plasmid (courtesy of the UF CTRND) using Lipofectamine LTX Reagent with PLUS Reagent (Invitrogen 15338030) according to its included protocol. Cells were maintained under hygromycin to only select for cells that were successfully transfected.

### Total Protein Extraction/Solubilization

MEFs cell lysates were prepared using a RIPA lysis buffer (1% TritonX, 50 mM Tris-HCl, 150 mM NaCl, 0.1% SDS) containing protease and phosphatase inhibitor tablets and then centrifuged for 15 minutes at 1400g at 4° C. Protein concentration was determined using the Pierce BCA protein assay kit (Pierce 23225). The protein sample was mixed with 4x laemmli buffer (Bio-Rad 1610747) and boiled on a heat block at 95° C for five minutes.

### Western Blot Detection of Progranulin, LAMP1 and Cathepsin B

Proteins were resolved by Mini-Protean TGX stain-free gels 4-20% (BioRad 4568093) and transferred to TransBlot Turbo mini-size PVDF membrane (BioRad). Membranes were fixed with 0.4% paraformaldehyde (PFA) then rinsed three times with ultrapure water using the Milli-Q EQ 7000 Ultrapure Water Purification System. The membrane was blocked in 5% powdered milk in 1X tris-buffered saline with Tween-20 (TBS-T), and primary antibodies and secondary antibodies were dissolved in this same buffer. Information on the antibodies used can be found in Table 1. Primary antibodies used were LAMP1 (rabbit, Abcam ab24170) diluted 1:1000, PGRN (sheep, R&D AF2557) diluted 1:400 and Cathepsin B (CTS B)(rabbit, Abcam ab214428) diluted 1:1000. Secondary antibodies Goat anti Rabbit HRP (Jackson Immunoresearch 111-035-144) and Donkey anti-Sheep HRP (Invitrogen A16041) were diluted 1:2000. The membrane was developed using SuperSignal West Pico PLUS Chemiluminescent Substrate (Thermo scientific 34577) and SuperSignal West Femto Maximum Sensitivity Substrate (Thermo scientific 34096) and visualized on the Li-cor Odyssey FX system.

### Immunocytochemistry

MEFs were plated in a 12-well plate at a density of 15,000 cells per well on top of Poly-D-Lysine/Laminin Cellware 12-mm round coverslips (Corning 354087) and incubated at 37° C overnight. The BMV109 probe was administered the next day and cells were fixed with 4% paraformaldehyde (PFA). MEFs were permeabilized with permeabilization buffer containing.05% TritonX, 1% normal donkey serum, and 1X TBS. They were blocked with a buffer containing 1% normal donkey serum and 1X TBS. Information of antibodies can be found in Table 2. LAMP1 (rat, Developmental Studies Hybridoma Bank 1D4B) and Tubulin-□ (mouse, Calbiochem cp06) primary antibodies were diluted 1:200 and 1:500 respectively in TBS plus 1% normal donkey serum and 0.1% Triton X-100. Donkey anti-rat AF488 (Invitrogen A48269) and donkey anti-mouse AF568 (Invitrogen A10037) secondary antibodies were diluted 1:2000 in TBS plus 1% normal donkey serum and 1% Triton X-100. MEFs were counterstained with DAPI (Invitrogen D13306) diluted 1:1000 in TBS. Prolong glass antifade mounting solution (Invitrogen P36982) was used prior to coverslipping.

### Statistics

Unpaired t-test analyses were performed on western blot data for LAMP1 expression. One-way ANOVA analyses were performed on western blot data for pan-cathepsin activity and all of the data from the N-TAP progranulin rescue experiments.

## Results

### Baseline expression of PGRN in Grn+/+ and Grn-/-MEFs

The first step of this study was to confirm that the model was lacking PGRN protein by western blot. It was confirmed that the *Grn*^*-/-*^ MEFs did not express PGRN and the *Grn*^*+/+*^ MEFs did (Fig. 1) which was consistent with the predicted PGRN expression profile of this model.

**Figure 1.**
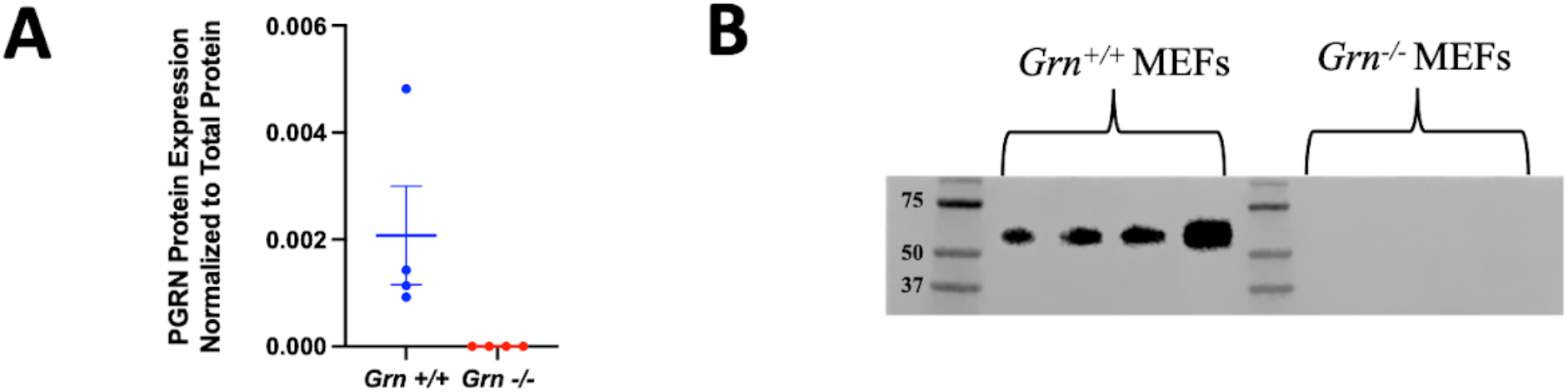
Baseline expression of PGRN in Grn+/+ and Grn-/-MEFs. (A) PGRN protein levels were quantified by western blot. (B) Representative western blot detection of PGRN protein in Grn+/+ MEFs and Grn-/-MEFs.

### Loss of progranulin results in increased pan-cathepsin activity in MEFs

After confirming the presence (*Grn*^*+/+*^) or absence (*Grn*^*-/-*^) of PGRN in the respective MEFs, pan-cathepsin activity was measured using the BMV109 pan-cathepsin probe. Using western blots, it was found that *Grn*^*-/-*^ MEFs had higher cathepsin activity than *Grn*^*+/+*^ MEFs (Fig. 2B). A one-way ANOVA analysis showed that the difference in pan-cathepsin activity between *Grn*^*-/-*^ MEFs and *Grn*^*+/+*^ MEFs is statistically significant, with a p-value of less than 0.01 (Fig. 2A). Next, this study attempted to confirm this difference in cathepsin activity between *Grn*^*-/-*^ MEFs and *Grn*^*+/+*^ MEFs using immunocytochemistry. An analysis of the integrated intensity of the BMV109 signal per cell revealed a trend towards increased pan-cathepsin activity in *Grn*^*-/-*^ MEFs (Fig. 2E). To see if the increased pan-cathepsin expression in the *Grn*^*-/-*^ MEFs was just a result of increased lysosomal health, lysosomal marker LAMP1 protein expression was measured by western blot. Using an unpaired t-test, it was found that there is a statistically significant increase in LAMP1 expression in *Grn*^*+/+*^ MEFs as compared to *Grn*^*-/-*^ MEFs (Fig. 2B). Figure 2 is a representative sample of multiple independent experiments.

**Figure 2.**
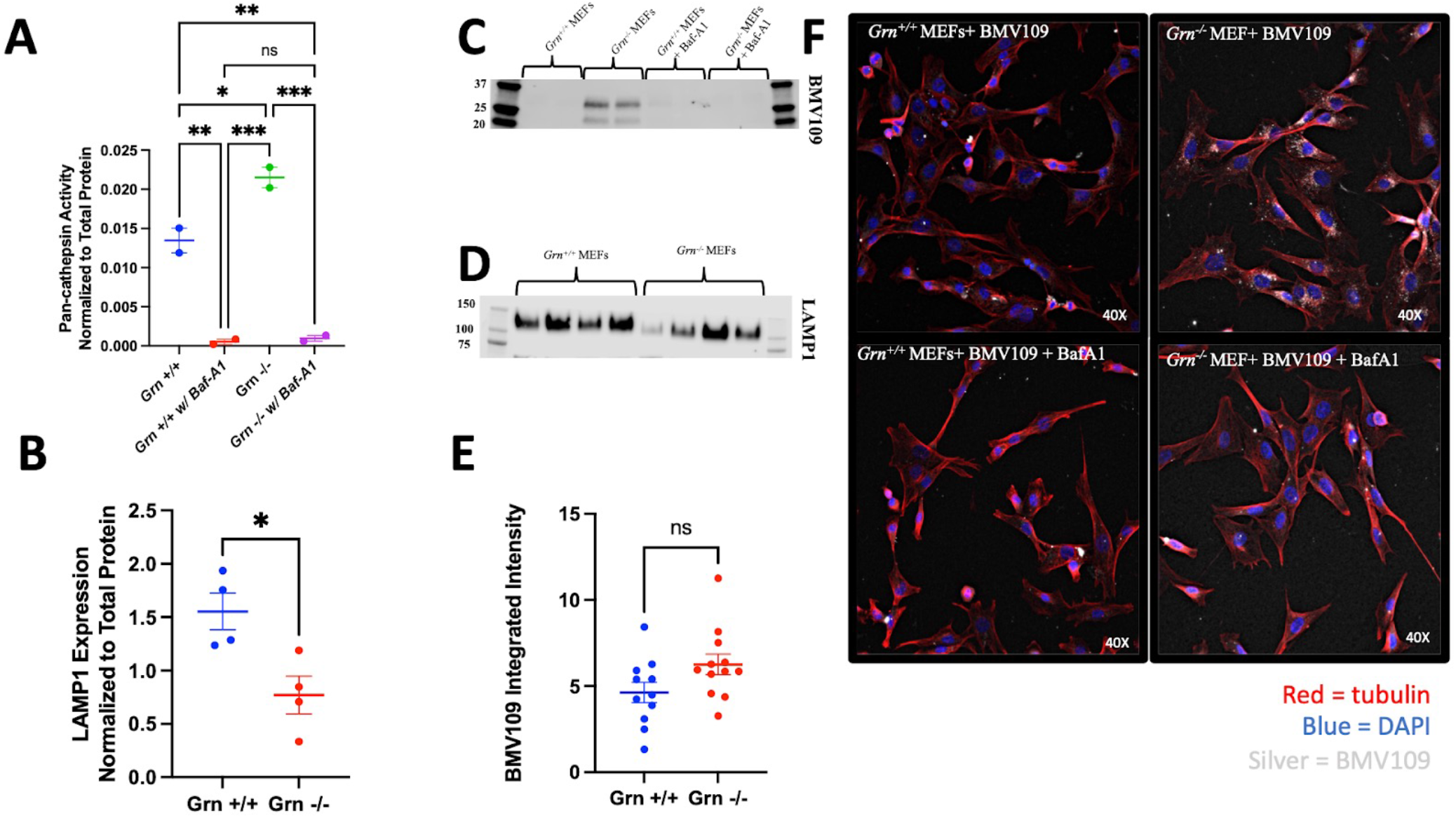
Loss of progranulin results in increased pan-cathepsin activity in MEFs. (A) Pan-cathepsin activity was quantified by western blot with BMV109. (B) LAMP1 expression was measured by western blot. (C) Representative western blot of BMV109. (D) Representative western blot of LAMP1. (E) Integrated intensity of BMV109 signal measured by immunocytochemistry. (F) Representative immunocytochemical detection of BMV109 in MEFs.

### N-TAP progranulin rescues pan-cathepsin activity and LAMP1 levels in Grn-/-MEFs

This stage of the investigation is aimed to partially restore PGRN protein expression in *Grn*^*-/-*^ MEFs to assess how pan-cathepsin activity and lysosomal health are impacted. To evaluate the impact on PGRN recovery in *Grn*^*-/-*^ MEFs, the cells were transfected with a plasmid expressing N-terminal tandem affinity purification (N-TAP) tagged PGRN. The results so far show no change in PGRN expression between *Grn*^*-/-*^ MEFs and *Grn*^*-/-*^ MEFs + NTAP PGRN (Fig. 3A). For pan-cathepsin activity, there is no significant difference between *Grn*^*+/+*^ MEFs, *Grn*^*-/-*^ MEFs and *Grn*^*-/-*^ MEFs + NTAP PGRN although there does appear to be a trend in the pan-cathepsin activity in the *Grn*^*-/-*^ MEFs + NTAP PGRN being reduced in comparison to the *Grn*^*-/-*^ MEFs so more experiments will be required to determine the significance of the findings. For LAMP1 expression, again there is no significant difference between LAMP1 expression in between *Grn*^*+/+*^ MEFs, *Grn*^*-/-*^ MEFs and *Grn*^*-/-*^ MEFs + NTAP PGRN but there is a trend that LAMP1 expression is increased in *Grn*^*-/-*^ MEFs + NTAP PGRN so more experiments will be required to determine the significance of the findings.

**Figure 3.**
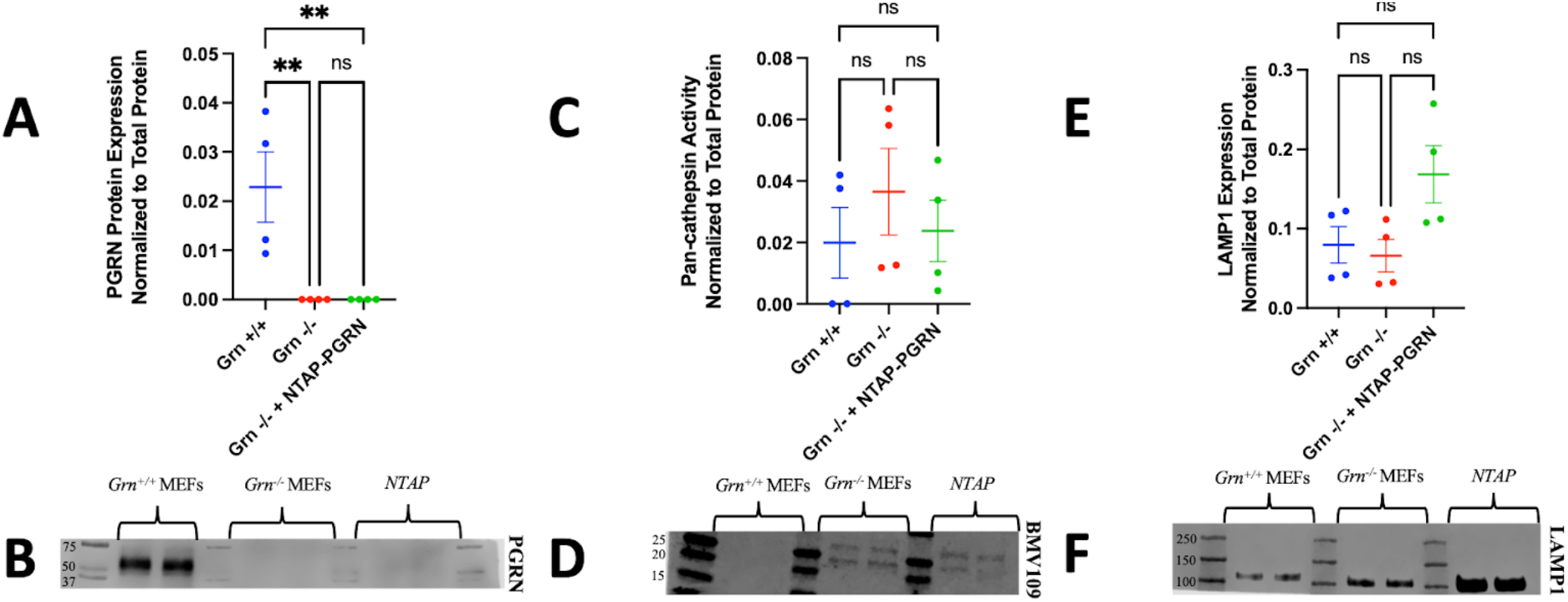
N-TAP progranulin rescues pan-cathepsin activity and LAMP1 levels in Grn-/-MEFs. (A) PGRN protein expression was quantified by western blot. (B) Representative western blot of PGRN protein. (C) Pan-cathepsin activity was determined by western blot with BMV109. (D) Representative western blot of BMV109. (E) LAMP1 expression was determined by western blot. (F) Representative western blot of LAMP1.

### Cathepsin B activity is increased in Grn-/-MEFs and rescued by N-TAP progranulin

After assessing pan-cathepsin activity using the BMV109 probe, to get more specific, an aim of this study is to use specific cathepsin antibodies to identify which exact cathepsins are having variable levels of activity between *Grn*^*-/-*^ MEFs and *Grn*^*+/+*^ MEFs. Although there are many cathepsins, this study has started with cathepsin B because it PGRN protein processing (Mohan et al., 2021). A one-way ANOVA analysis showed that there is a statistically significant difference in cathepsin B activity between *Grn*^*-/-*^ MEFs and *Grn*^*+/+*^ MEFs (Fig. 4A) with a p-value of less than.01. There is also a statistically significant difference in cathepsin B activity between *Grn*^*-/-*^ MEFs and *Grn*^*-/-*^ MEFs + NTAP PGRN (Fig. 5A) with a p-value of less than.05. However, there is not statistically significant difference in cathepsin B activity between *Grn*^*+/+*^ MEFs and *Grn*^*-/-*^ MEFs + NTAP PGRN (Fig. 4A). Figure 4 is a representative sample of multiple independent experiments.

**Figure 4.**
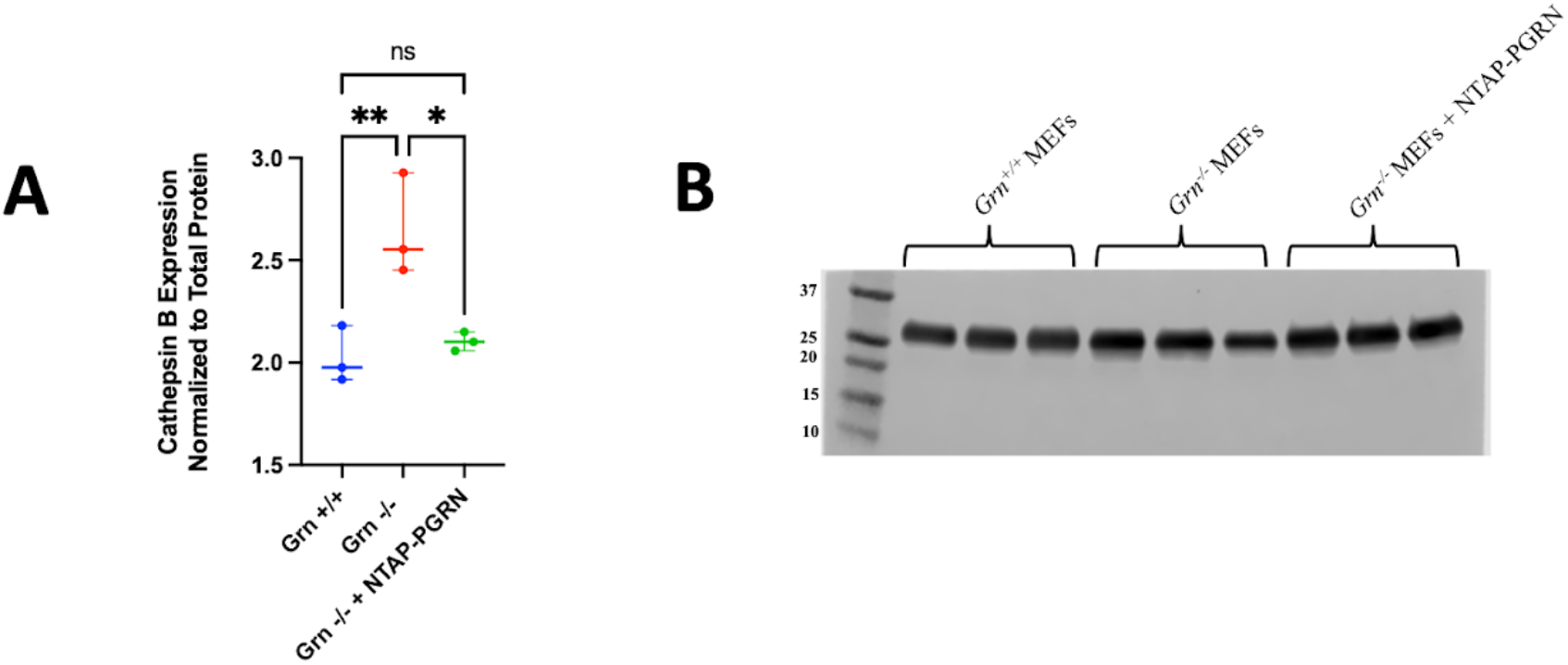
Cathepsin B activity is increased in Grn-/-MEFs and rescued by N-TAP progranulin. (A) Cathepsin B activity was measured by western blot. (B) Representative western blot of cathepsin B in MEFs.

## Discussion

These studies revealed that there is a statistically significant difference in pan-cathepsin protein expression between *Grn*^*-/-*^ MEFs and *Grn*^*+/+*^ MEFs, as measured by western blot analysis and confirmed by immunocytochemistry. It was shown that *Grn*^*-/-*^ MEFs have higher pan-cathepsin expression than *Grn*^*+/+*^ MEFs. Because cathepsins are lysosomal proteases, it was important to confirm that the statistically significant difference in pan-cathepsin activity between *Grn*^*-/-*^ MEFs and *Grn*^*+/+*^ MEFs was not just a result of one genotype having better lysosomal health than the other. Western blots demonstrated that *Grn*^*+/+*^ MEFs displayed significantly more LAMP1 expression than *Grn*^*-/-*^ MEFs, confirming that the increase in pan-cathepsin activity in *Grn*^*-/-*^ MEFs was not just a result of increased lysosomal health in that genotype. Drawing the conclusion that *Grn*^*-/-*^ MEFs have increased cathepsin activity has implications in the context of inflammation. More specifically, this study demonstrates that there is increased cathepsin B activity in *Grn*^*-/-*^ MEFs in comparison to *Grn*^*+/+*^ MEFs and this cathepsin B activity is reduced when *Grn*^*-/-*^ MEFs are rescued with N-TAP PGRN. As discussed in the introduction, cathepsins play an important role in triggering an immune response which includes inflammation. One possible explanation for this is that there is increased cathepsin activity in the *Grn*^*-/-*^ MEFs because the cathepsins are acting as a compensatory mechanism to make up for the loss of PGRN protein to help the lysosome to continue to perform its normal functions. To investigate this directly, there are ongoing studies with the N-TAP PGRN cell line to determine the extent to which PGRN rescue changes pan-cathepsin activity in the MEFs. It would also be beneficial as a future direction to investigate cathepsin activity in cells that express PGRN protein in large amounts such as microglia which are directly related to inflammation and neurodegeneration.

One limitation of these studies is the use of the MEF model. There are advantages to using MEFs for studying biochemical processes such as their rapid growth rate, however there are also limitations (Qiu et al., 2016). This model does not serve as the most representative model of what is happening in the brain so it would be beneficial to repeat the assays in primary cells such as microglia from *Grn*^*-/-*^ mice and *Grn*^*+/+*^ mice because PGRN protein is highly expressed in microglia (Götzl et al., 2018). Another limitation of this study is BMV109, the pan-cathepsin probe used to assess cathepsin activity. It is noticeable in the blot found in Figure 3C that the signal from BMV109 appears as multiple bands that look slightly smeared. It is possible that the blots appear this way because multiple different cathepsins are being detected, and different cathepsins appear at different molecular weights. Future directions should also include investigating different specific cathepsins beyond cathepsin B, such as cathepsin L and cathepsin D.

## Author contributions

AA designed, executed, and analyzed the experiments. She wrote the manuscript under supervision of MGT. All authors edited the manuscript.

## Acknowledgements

We thank Cody Keating and the Tansey lab for useful discussions. We thank Andy Nguyen at SLU (and Robert V. Farese) for providing the Grn-/- and Grn+/+ MEFs. This work was partially funded by a University of Florida University Scholars Program (to AA), the Summer Neuroscience Institute Program (SNIP) at the University of Florida in the Department of Neuroscience (to AA), and NIH/NINDS grant RF1NS28800 (to MGT).

